# minicore: Fast scRNA-seq clustering with various distances

**DOI:** 10.1101/2021.03.24.436859

**Authors:** Daniel N. Baker, Nathan Dyjack, Vladimir Braverman, Stephanie C. Hicks, Ben Langmead

**Affiliations:** Department of Computer Science, Johns Hopkins University; Department of Biostatistics, Johns Hopkins Bloomberg School of Public Health

## Abstract

Single-cell RNA-sequencing (scRNA-seq) analyses typically begin by clustering a gene-by-cell expression matrix to empirically define groups of cells with similar expression profiles. We describe new methods and a new open source library, `minicore`, for efficient *k*-means++ center finding and *k*-means clustering of scRNA-seq data. `Minicore` works with sparse count data, as it emerges from typical scRNA-seq experiments, as well as with dense data from after dimensionality reduction. `Minicore`’s novel vectorized weighted reservoir sampling algorithm allows it to find initial *k*-means++ centers for a 4-million cell dataset in 1.5 minutes using 20 threads. `Minicore` can cluster using Euclidean distance, but also supports a wider class of measures like Jensen-Shannon Divergence, Kullback-Leibler Divergence, and the Bhattacharyya distance, which can be directly applied to count data and probability distributions.

Further, `minicore` produces lower-cost centerings more efficiently than scikit-learn for scRNA-seq datasets with millions of cells. With careful handling of priors, `minicore` implements these distance measures with only minor (*<*2-fold) speed differences among all distances. We show that a `minicore` pipeline consisting of *k*-means++, localsearch++ and minibatch *k*-means can cluster a 4-million cell dataset in minutes, using less than 10GiB of RAM. This memory-efficiency enables atlas-scale clustering on laptops and other commodity hardware. Finally, we report findings on which distance measures give clusterings that are most consistent with known cell type labels.

**Availability:** The open source library is at https://github.com/dnbaker/minicore. Code used for experiments is at https://github.com/dnbaker/minicore-experiments.

## 1 Introduction

Single-cell RNA-sequencing (scRNA-seq) is capable of measuring transcriptome-wide gene expression in millions of cells per experiment. With the arrival of multi-million-cell datasets [9, 12], and larger efforts like the Human Cell Atlas [24] on the horizon, the need for methods that rapidly analyze and cluster (empirically group) cells is growing. This necessitates computational advances in methods for unsupervised clustering and summarizing large collections of cells.

*k*-means is one popular clustering framework. It is classically formulated as an expectation maximization problem that starts from an initial set of *k* data points that act as “centers” [20], iterating to obtain final centers. These centers induce a clustering of the observations into *k* classes. *k*-means++ [2] improves how the initial centers are found, yielding clear mathematical guarantees for the overall clustering. Besides its direct application as clustering methods, *k*-means and *k*-means++ are useful as individual components of other methods, including for data quantization [20], spectral clustering [10], outlier detection [28], machine learning [1] and construction of sketches and coresets [22, 14]. For example, in scRNA-seq analysis, sketching – the selection of a possibly weighted subset of cells to use – can be used to identify rare cell types. The Geometric sketching [16], Hopper [13], and submodular sketch [30] methods all employ some form of center-finding as a subroutine.

We describe a new open source, highly efficient library software library called `minicore`, which implements an array of algorithms to find the “center” of a group of cells – essentially a rough clustering – and for performing *k*-means clustering seeded by those centers. The advantages of `minicore` are threefold. First, `minicore` uses a new vectorized weighted reservoir sampling algorithm for its initial center-finding step, making it far more efficient than competing *k*-means++ algorithms like that in scikit-learn. Second, `Minicore` implements a variety of distance measures, including the widely-used squared Euclidean distance, but also including others like Jensen-Shannon Divergence, Kullback-Leibler Divergence, and Bhattacharyya distance,, which can be directly applied to count data or probability distributions. Third, `minicore` is able to process both dense, dimensionality-reduced data – the typical input for scRNA-seq clustering methods – as well as full, sparse, non-reduced matrices of counts. `Minicore` is unique in its ability to handle scRNA-seq data in both sparse and dense forms, and its support for distance measures that account for the original count-based nature of the data.

On real scRNA-seq datasets with up to millions of cells and using squared Euclidean distance, `minicore` is substantially faster than scikit-learn and achieves lower objective-function cost. Further, `minicore` can produce centers using a wide variety of distance measures with only minor differences in the overall running time, facilitating use of distance measures that are better attuned to the count nature of the data and do not require prior transformations [27]. Finally, we show that a complete pipeline consisting of `minicore`’s implementations of *k*-means++, localsearch++ and mini-batch *k*-means can cluster a 4-million cell dataset in minutes using 20 threads and a maximum resident set size (RAM) of less than 10 GiB.

## 2 Results

We collected scRNA-seq datasets of varying size: (a) the PBMC dataset consisting of 68,579 peripheral blood mononuclear cells (PBMC) from human [32], (b) the Cao et al mouse organogenesis dataset (Cao2m) consisting of 2,058,652 cells [9], and (c) the Cao et al human fetal gene expression dataset (Cao4m) consisting of 4,062,980 cells [8]. In all cases, the original form of the data is a sparse matrix of gene-by-cell nonnegative integer counts. For datasets not originally represented in compressed-sparse-row (CSR) format, we convert them to that format prior to our experiments. Each of the three datasets has an associated set of cell-type labels, obtained by the original authors through an analysis that combined an initial clustering with foreknowledge of specific marker genes [32, 9, 8]. While these label assignments are not “ground truth,” they capture some biological foreknowledge and so we use them to evaluate our final clusterings below.

While `minicore` can cluster sparse counts directly, we also generated a dense version of each of the three datasets after applying a dimensionality reduction method. Specifically, we used the truncated Singular Value Decomposition (SVD) from scikit-learn. Rows of the final matrix consist of the original data’s projection into the first 500 principal components. We note that a standard PCA has a “centering” step where the mean is subtracted from each feature. We used a non-centered SVD since centering causes the matrix to lose its zero entries and become dense, in turn requiring would require terabytes of memory for an SVD computation over millions of cells. While non-centered SVD avoids this problem by keeping the matrix sparse, a drawback is that the resulting principal components are selected based not just on the amount of variability but also on the magnitudes of the values. This is addressed further in Discussion.

### 2.1 Fast and accurate center finding with minicore *k*-means++

We used `minicore` v0.3 and compared it to scikit-learn’s v0.12.4 function for *k*-means++ center finding (`sklearn.cluster.kmeans plusplus`). We considered various values for the number of centers, *k*. We note that scikit-learn supports only the squared Euclidean distance measure and does not support the use of multiple threads in parallel. For the most direct comparison, we used a single thread and the squared Euclidean distance only. In all cases, we measured the running time and squared-Euclidean objective cost of the resulting set of centers. In the case of `minicore`, we benchmarked both the *k*-means++ method (MC), as well as the *k*-means++ method augmented by localsearch++ (MCLS). The scikit-learn results are labeled SKL.

Using the three datasets, we found that our `minicore` *k*-means++ (MC) implementation is significantly faster when compared to scikit-learn *k*-means++ (SKL) using both sparse and dense data (Figure 1, Table 1). For dense input data, the MC mode of `minicore` had a dramatic speed advantage, achieving 100–150 times greater speed for the PBMC dataset compared to scikit-learn, about 50–100 times greater for Cao2m, and about 240–280 times greater for Cao4m. For sparse data, the MC mode of `minicore` was 3–9 times faster than scikit-learn depending on the experiment. Our implementation of `minicore` *k*-means++ augmented by the localsearch++ procedure (MCLS) was also always faster than SKL, and was only about 2.5–5 times slower than the MC mode, depending on the experiment.

**Table 1:**
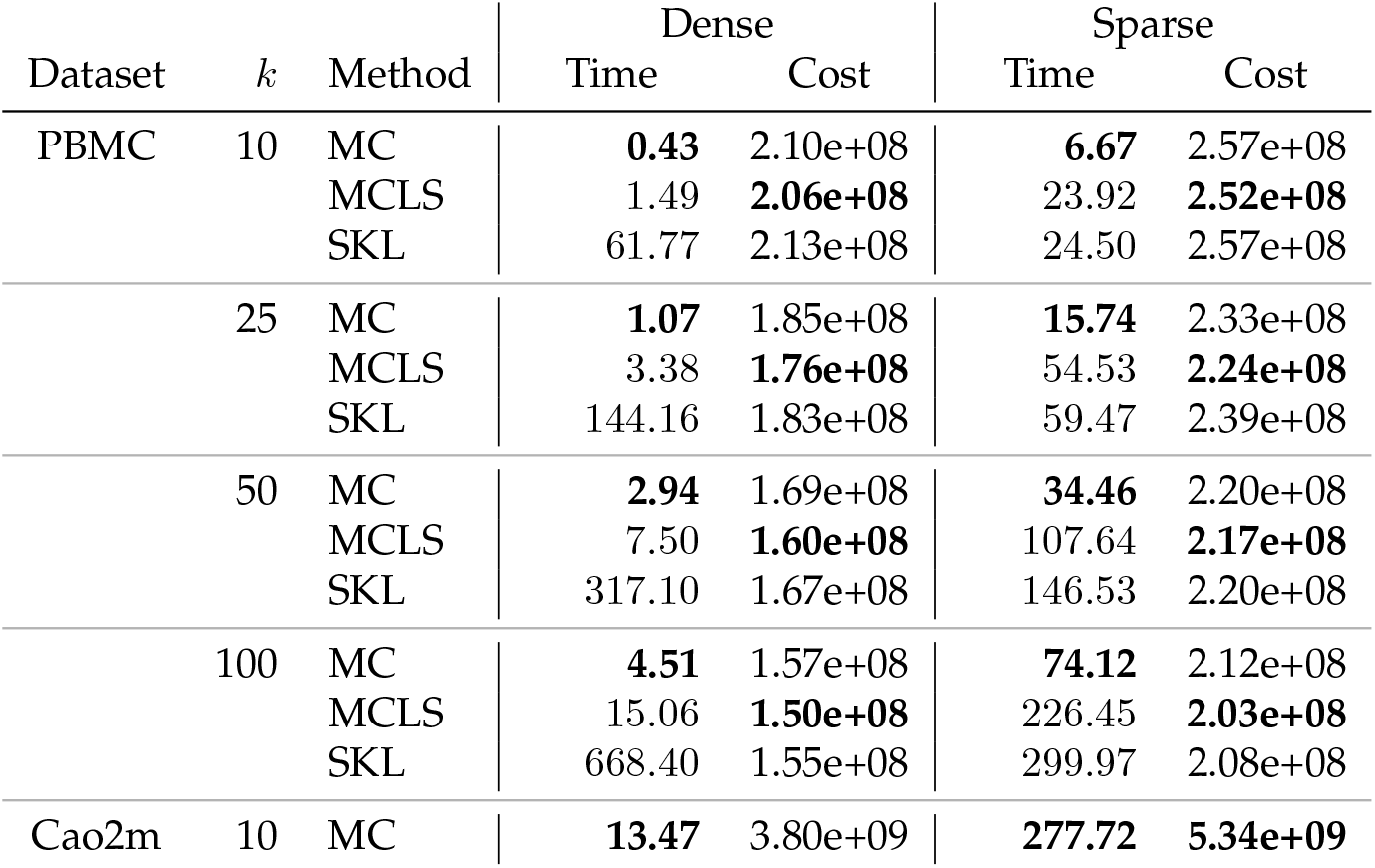
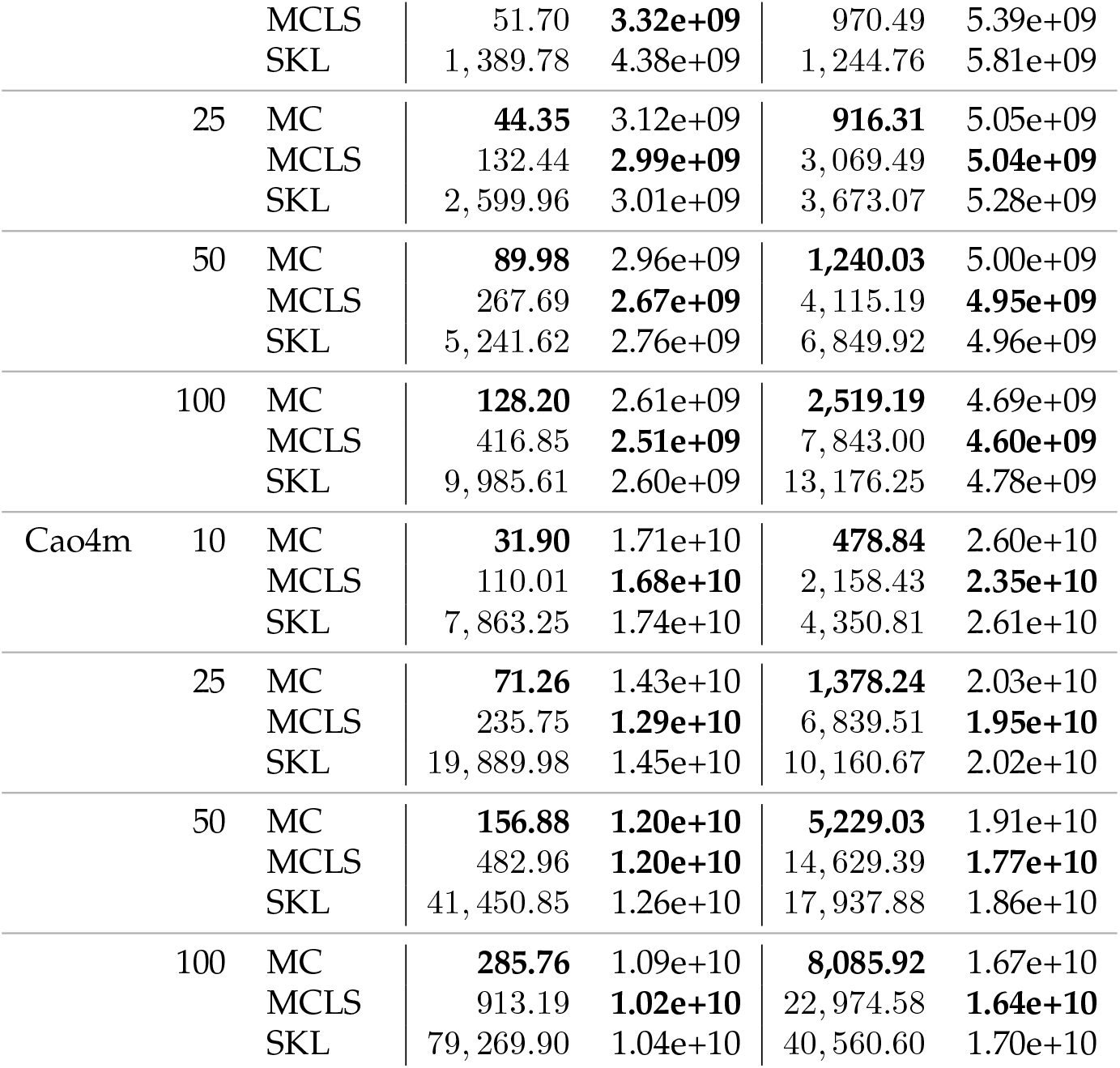
*k*-means++ clustering results for `minicore` (MC), `minicore` with localsearch++ (MCLS), and scikit-learn (SKL). All experiments use squared Euclidean distance and a single thread of execution.

**Figure 1.**
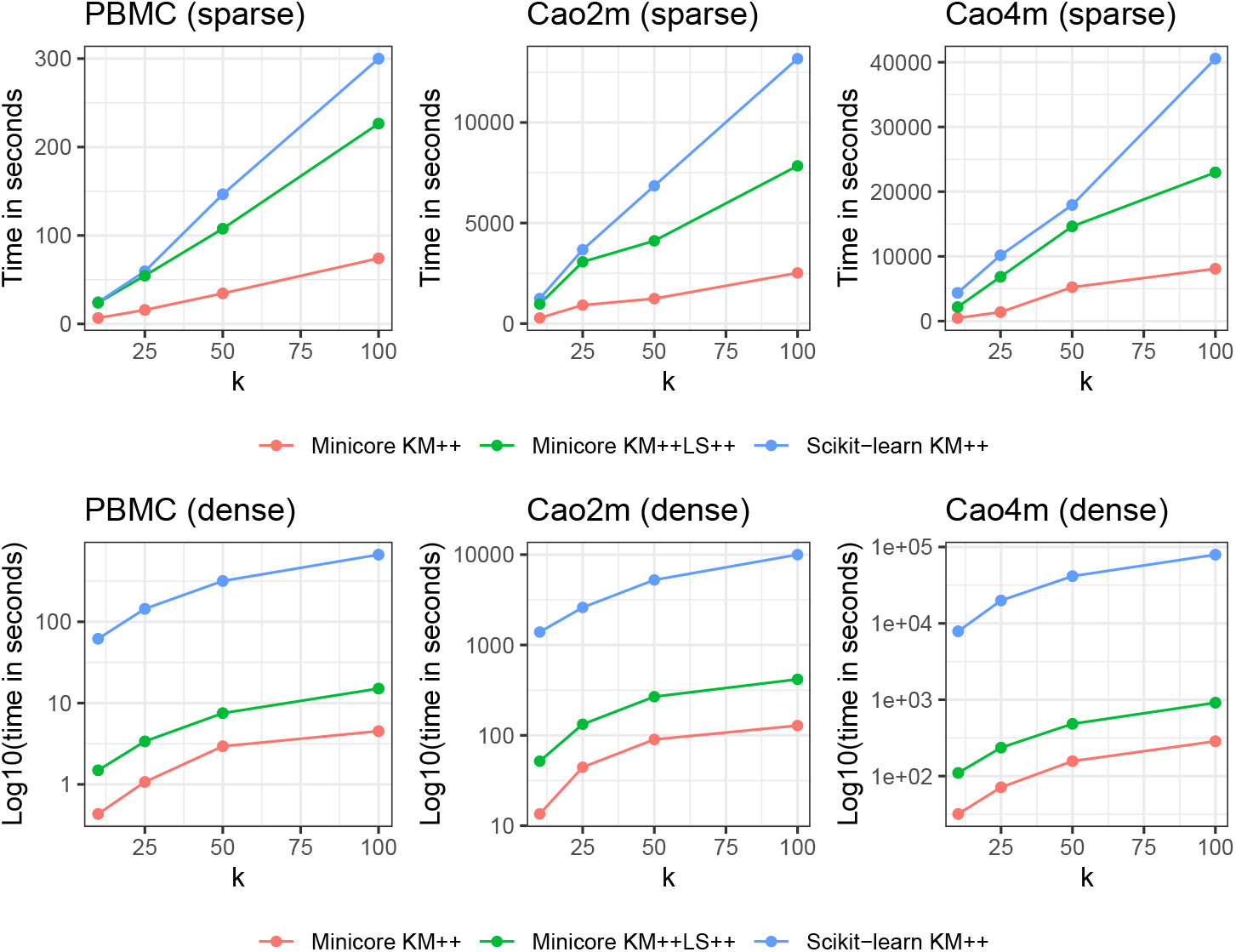
minicore *k*-means++ is faster than scikit-learn *k*-means++. Performance evaluation (y-axis) of elapsed time (seconds) for sparse data (top) and log_10_ transformed time for dense data (bottom) for increasing sizes of *k* (x-axis) for the PBMC dataset with 68k cells (left), Cao et al. dataset with 2 million cells (middle), and Cao et al. dataset with 4 million cells (right). Results for `minicore` *k*-means++ are in red (standard) and green (with localsearch++); scikit-learn *k*-means++ is blue.

Further, we found that both of our `minicore` *k*-means++ implementations (MC and MCLS) obtained comparable or lower costs of the objective function compared to SKL (Table 1). MCLS obtained the lowest objective in nearly all cases across the three datasets (both dense and sparse).

Overall, the results showed that `minicore` produces high-quality centers and readily scales to multi-million cell datasets, even in their original sparse form. For example, the MC mode used about 2h:15m (single-threaded) to find *k* = 100 centers for the 4-million cell Cao4m dataset.

Similarly, `minicore` makes economical use of memory even when working directly on sparse representations. The 4-million cell dataset can be clustered using less than 10 GiB RAM, allowing it to run on commodity hardware.

### 2.2 minicore supports distance metrics for both continuous and count data, and probability distributions

To evaluate `minicore`’s speed for distance measures beyond the commonly used squared Euclidean distance (SQE), we ran `minicore` using other measures, including the Bhattacharyya Metric (BAT), Kullback-Leibler Divergence (KLD), Jensen-Shannon Divergence (JSD), and cosine distance (COS). While these measures involve computationally demanding operations like logarithms and square roots, `minicore` optimizes these inner loops using the SLEEF library and vectorization [26]. An additional challenge is the need to handle 0 counts, which can infinite divergence for measures like the KLD. To address this, we use a lazily applied prior that avoids having to instantiate a dense version of the matrix at any point. See Methods for more details on both these points.

Using the 2 million and 4 million Cao et al. datasets, we found that the choice of distance metric used for `minicore`’s *k*-means++ algorithm does impact speed, but not dramatically (Figure 2). Specifically, we found that the Bhattacharyya Metric (BAT) required less time than squared Euclidean in all cases, whereas KLD required roughly the same amount of time as SQE, and JSD generally required the most time. Importantly, the slowest measure (often the JSD) never requires more than 61% more computation time than the fastest measure.

**Figure 2.**
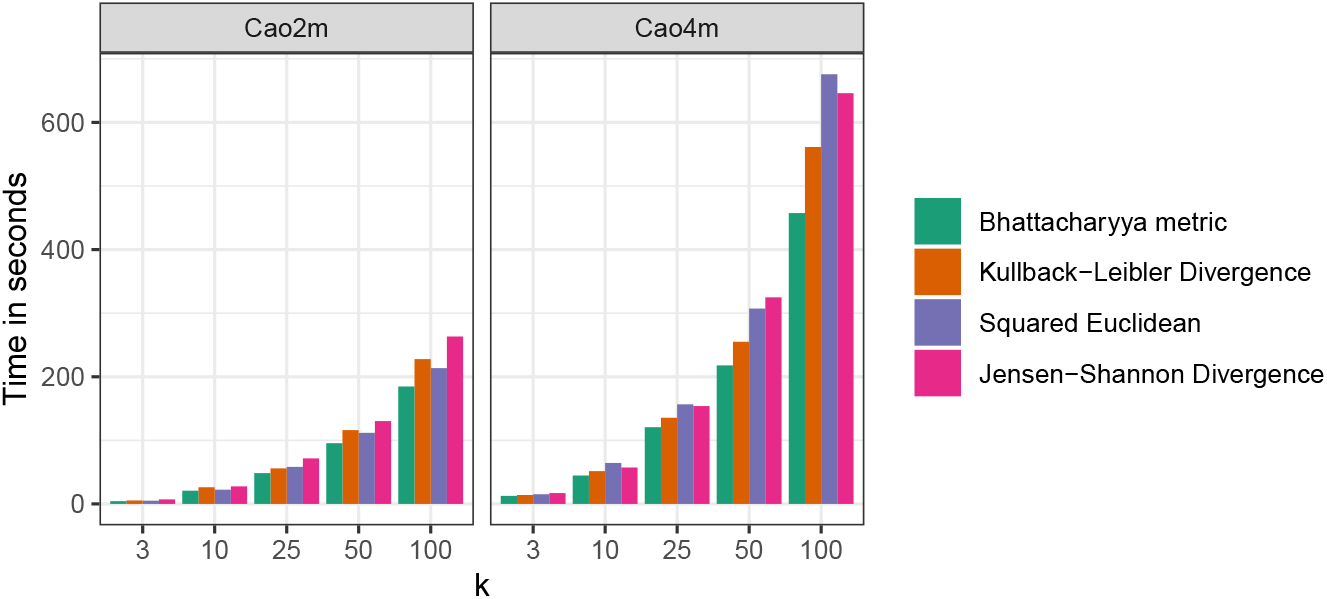
The choice of distance has minor impact on the speed of minicore *k*-means++. Performance evaluation (y-axis) of elapsed time (seconds) for sparse data for increasing sizes of *k* (x-axis) for Cao et al. dataset with 2 million cells (left), and Cao et al. dataset with 4 million cells (right). For a given dataset and *k*, the slowest measure never requires more than 61% more time than is required by the fastest measure. All experiments used 16 simultaneous threads and the localsearch++ improvement was not run.

### 2.3 minicore supports *k*-means and mini-batch *k*-means clustering algorithms

The `minicore` library also supports both full *k*-means clustering using Lloyd’s algorithm [21], and the faster mini-batch *k*-means algorithm [25, 15]. We sought to measure the efficiency and accuracy of a full *k*-means clustering pipeline built from the *k*-means++, localsearch++, and mini-batch *k*-means components of the `minicore` library. We chose mini-batch *k*-means rather than Lloyd’s algorithm because the mini-batch approach has recently been shown to be significantly faster for large datasets and provides similar results [15]. In all cases, we used *k* = 25, a mini-batch *k*-means batch size of 10,000, 25 rounds of localsearch++, and a prior of 0.01

We again analyzed the PBMC, Cao2m and Cao4m datasets. We evaluated the clusterings using the cell-type labels provided by the authors of the datasets [32, 9, 8]. Specifically, we used our *k*-means clusters as empirical cell labels, comparing these to the “true” labels using the Adjusted Rand Index (ARI).

While we began with the full sparse matrix, we subsampled the rows to consist of the 500 most variable genes [7], as this often achieved greater Adjusted Rand Index compared to analyzing the entire matrix. We tried several distance measures: Bhattacharyya Metric (BATMET), Jensen-Shannon Divergence (JSD), the Kullback-Leibler Divergence (KLD), and Squared Euclidean Distance (SQE). We ran `minicore` using 20 simultaneous threads.

We found that `minicore` was able to cluster the cells in all three datasets in minutes, with the slowest experiment taking about 12 minutes (Figure 3). For the Cao2m and Cao4m datasets, timings were in the range of 325–365 seconds and 200–700 seconds respectively.

**Figure 3.**
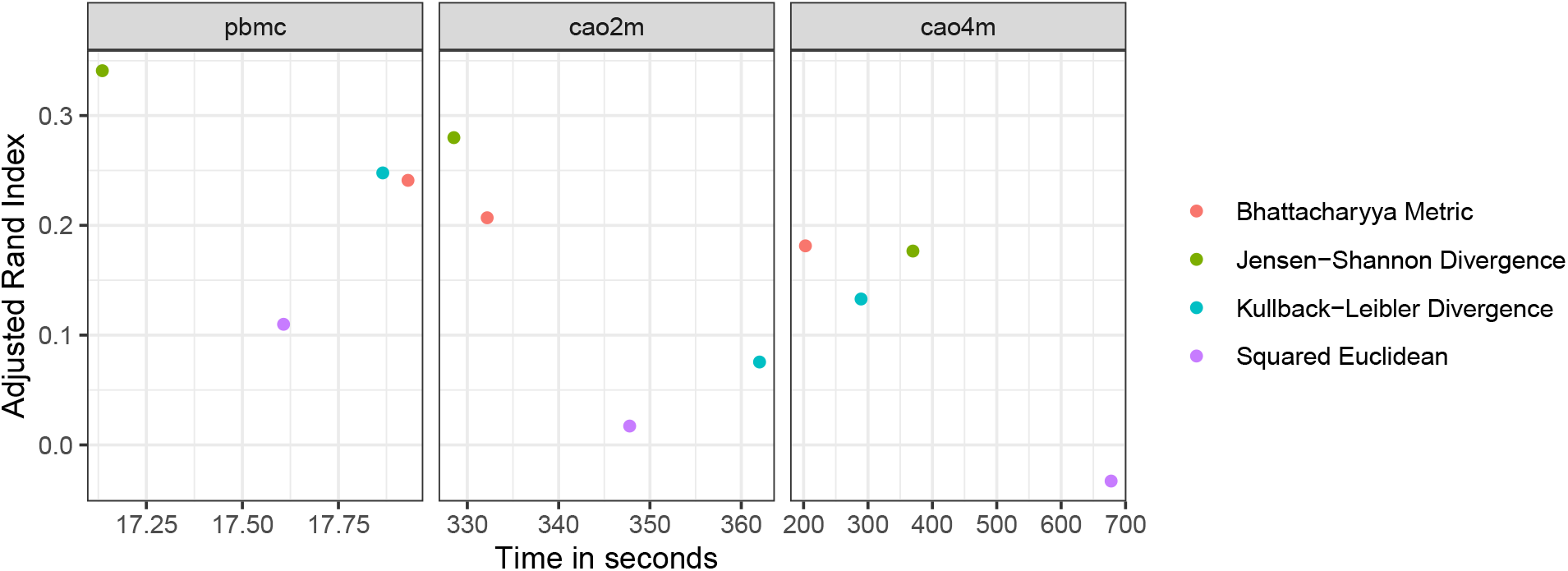
Clustering accuracy (ARI, vertical) versus running time (seconds, horizontal) for various datasets and distance measures. All experiments used the 500 most variable genes, *k* = 25, a mini-batch *k*-means batch size of 10,000, 25 rounds of localsearch++, and a prior of 0.01.

For both Cao2m and Cao4m, the Bhattacharyya Metric (BATMET) was superior to the Kullback-Leibler Divergence (KLD) and Squared Euclidean distance (SQE), achieving both greater speed and a higher Adjusted Rand Index for its final clustering. In the case of Cao2m, the JSD was superior to BATMET on both speed and ARI, but this relationship is reversed for the Cao4m dataset.

We measured `minicore`’s peak memory footprint (resident set size) when processing the Cao4m dataset and found that it was less than 10GiB RAM. In short, we found that `minicore` was capable of analyzing a 4-million cell dataset in a few minutes using computational resources consistent with a typical commodity laptop.

## 3 Discussion

We introduced a new library called `minicore` for *k*-means clustering of scRNA-seq datasets. An efficient, vectorized sampling kernel fuels both its *k*-means++ center finding algorithm and its localsearch++ algorithm for refining centers. Combined with an efficient mini-batch *k*-means implementation, these components form a complete and efficient pipeline for *k*-means clustering of scRNA-seq data, requiring about 3.5 minutes to cluster a *>*4 million cell dataset when using 20 threads and less than 10GiB RAM. This low memory requirement brings even atlas-scale clustering within reach of laptops and other commodity hardware. While we applied `minicore` to scRNA-seq here, its algorithms are readily adaptable to other applications, for instance in data quantization, outlier detection and spectral clustering [20, 10, 28].

`Minicore`’s fast implementations of various distance measures, gives users the flexibility to tailor the distance measure to the data. Different measures might be appropriate depending on whether cells are best viewed as vectors of real numbers, vectors of counts, or probability distributions. We showed that distance measures other than squared Euclidean can perform substantially better when evaluated using given cell type labels. In future work, we plan to explore how `minicore` can be applied beyond to work, for example, with graph-induced metrics [4].

Another likely application of the algorithms in `minicore` is to build “sketches” of large single-cell data compendia. A sketch is a weighted subset of cells that effectively span the gene-expression space and – like centers – facilitate the identification of accurate predicted cluster labels down-stream. Sketching approaches have been applied to the problem of obtaining cluster labels that accurately capture empirical groupings of rare cell types [13, 16, 31].

Finally, we further seek to explore whether our optimized weighted sampling kernel may also be applicable in the mini-batch *k*-means algorithm, specifically for the importance sampling required to drive the gradient-descent version of mini-batch *k*-means [6] [23].

In some experiments descried here, we used a non-centered version of the truncated Singular Value Decomposition (SVD) to project datasets into their first 500 principal components. We avoided the mean centering in order to keep the data sparse in preparation for the SVD. This has the drawback that the truncated SVD was selecting components based not only on variability, but also on the magnitudes of the points. In the future, we would like to address this by implementing or otherwise integrating a sparse version of a centered SVD computation into `minicore`. This could become an optional first step allowing users to create smaller, dense representations.

## 4 Methods

### 4.1 *k*-means++ algorithms in minicore

*k*-means gives an efficient way to choose an initial set of centers in preparation for the more work-intensive *k*-means optimization procedure. Unlike the simple strategy of choosing centers uniformly at random, *k*-means++ guarantees that the objective achieved by the downstream *k*-means procedure will be within a multiplicative *O*(log *k*) factor of the optimal cost objective.

The *k*-means++ algorithm involves choosing one center per step across *k* steps. In the first step, a center is chosen from among the data points uniformly at random. In subsequent steps, a new center is chosen in a weighted random fashion, with the probability of selecting a given point being proportional to its cost, specifically the distance to the nearest already-selected center. The algorithm therefore is a weighted sampling procedure. We now describe in detail, as similar sampling procedures form the core of multiple components of `minicore`.

#### Sampling kernel

In a given step of *k*-means++, a simple sampling strategy would be to calculate the cost of each as-yet-unchosen data point (potential “center” gene) then draw a random variate from a multinomial distribution weighted by those costs. Computationally, this can be accomplished in four steps: first, calculate a cost for each point, next calculate a prefix sum over the array of all costs, next generate a uniform random variate in [0, *C*] where *C* is the total cost, then perform binary search over the prefix-sum array to identify the point corresponding to the random variate. While binary search is fast, the costs, and therefore the prefix sum, must be at least partially re-computed in each of the *k* steps. Further, the prefix sum computation has an inherent dependence structure that inhibits parallelization, though *O*(*n* log *n*)-time parallel solutions exist.

`Minicore` instead uses a parallelized reservoir-sampling approach that extends an algorithm by Hü bschle-Schneider & Sanders [17]. That algorithm uses the fact that weighted sampling without replacement is equivalent to generating an exponential random variate for each data point, then selecting the point(s) with minimal variates. Importantly, variates can be drawn in parallel batches using single instruction multiple data (SIMD) instructions, providing instruction-level parallelism. Specifically, we use the SIMD-accelerated Polynomial Congruential Generator (PCG) SIMDPCG [19, 11]. Because variates are drawn independently for each point, `minicore` can additionally use multiple simultaneous threads to generate variates in parallel across processors.

While drawing the random variates involves a computationally expensive logarithm, we used the SLEEF library to compute batches of logarithms accurately and in parallel using SIMD instructions. As described in [17], exponential random variates can be sampled equivalently either by generated a random value *v* ∼*U* (0, 1) and exponentiating by the inverse of the weight 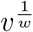 or, equivalently logging and dividing by the weight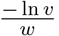 which is more numerically stable. We found this numerically stable alternative to be about 3 times as fast as exponentiating.

It is common for *k*-means++ implementations to select more than one potential new center in a single step, ultimately choosing the center that yields the lowest overall cost. Our parallel implementation accomplishes this using a per-thread heap data structure. SIMD instructions are used to determine which from among the random variates in a chunk are small enough to be added to the heap. If any are small enough, a serial loop extracts the variates and adds them. As a thread proceeds along the array of variates, heap updates become rarer, allowing the vast majority of the computation to remain SIMD parallelized. Finally, the samples in the per-thread heaps are combined to obtain an overall sample.

We can eliminate a significant number of branches in building the heap using Population Counts (popcount) and Count Trailing Zeros (CTZ) instructions. For each vector of new candidate variates, we compare it to the broadcasted ceiling, convert to a bitmask, and popcount, and switch on the value of the popcount, performing the heap update once per nonzero in the bitmask. We access the “current” bit by counting trailing zeros and indexing the relevant variate.

This sampling kernel is a core feature of our library, accessible with a C and C++ APIs in the free and MIT-licensed [3] library. While we described the sampling approach in the context of *k*-means++, it also forms the core of the localsearch++ algorithm described below.

#### localsearch++

Lattanzi and Sohler suggested an augmentation of *k*-means++ that adds sampling with local search heuristics [18]. At each iteration in localsearch++, there is an additional final step that selects the center that contributes the least to the objective. It eliminates that center and replaces it by a newly-sampled point. We re-use the previous sampling kernel to implement the weighted sampling required by this approach. To our knowledge, this is the first application of localsearch++ to distance measures beyond squared Euclidean distance.

### 4.2 Distance measures and sparsity in minicore

While *k*-means++ is most commonly implemented using Euclidean distance, it has also been shown that the *k*-means++ procedure yields a (log *k*)-approximate solution in expectation when using other distance measures and divergences [5]. Specifically, this applies to the class known as Bregman divergences, as well as convex combinations thereof. This class includes relevant measures such as Kullback-Leibler Divergence (KLD), Jensen-Shannon Divergence (JSD), Squared Euclidean distance (SQE) and others. Given this fact, we decided to implement the four distance measures detailed in Table 2.

**Table 2:** Formulas for distance measures implemented in `minicore`. Let *X*_*i*_, *Y*_*i*_ denote the *i*^*th*^observation (gene) for cells *X* and *Y*. Let 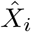 and *Ŷ_i_* denote the scaled (normalized by total cell-wide count) version of this entry. These all belong to the class of Bregman divergences, except the JSD which is a convex combination of Bregman divergences.

| Name | Abbreviation | Formula |
| --- | --- | --- |
| Squared Euclidean | SQE | $\sum_i (X_i - Y_i)^2$ |
| Kullbeck-Liebler Divergence | KLD | $\sum_i \hat{X}_i \times \log \frac{\hat{X}_i}{\hat{Y}_i}$ |
| Jensen-Shannon Divergence | JSD | $\frac{1}{2} \times (KL(X, \frac{X+Y}{2}) + KL(Y, \frac{X+Y}{2}))$ |
| Bhattacharyya Metric | BATMET | $\sqrt{1 - \sqrt{X} \cdot \sqrt{Y}}$ |

An important concern when implementing this other measures is how they handle 0 values in the data matrix. KL Divergences can be infinite for zero-valued entries, and other measures can have issues with numerical stability in these cases. This can be addressed by the use of a “prior” [29] of a Gamma(*β, β*) distribution, with a value *β >* 0. The *a posteriori* estimates are then *N*_*i*_ + *β*, ensuring no 0-valued entries. These are effectively “pseudo-counts,” a common way to adjust scRNA-seq data. Selecting *β* = 1 corresponds to a Dirichlet prior, while smaller values will penalize missing or low-count observations, and larger values will move points closer together for probability distribution-based distances.

This in turn creates another concern: a matrix adjusted by the prior will have no zero-valued entries, essentially becoming a dense matrix. This greatly increases the space and time required, making these distances impractical for large datasets. We instead compute distances with a lazy prior adjustment for all features, accounting for the zero-count features in aggregate. This is particularly advantageous for sparse matrices with a small number of nonzero values (*nnz*). In particular, we can perform distance computations in O(*nnz*) space and time rather than O(*d*), where *d* is the number of features. The general pseudocode for our distance computations is in Algorithm 1 1. For perspective, the 4-million cell dataset with 63,561 columns would require 960GiB of memory, nearly 100 times the 9.8GiB of the Compressed-Sparse Row (“CSR”) representation when using 16-bit indices and data fields. In this way, `minicore` can cluster atlas-scale datasets in reasonable working memory, operating directly on the sparse data.

### 4.3 Other optimizations

While minicore can cluster datasets in a fraction of the space the dense instantiation would require, it can scale even further while managing memory requirements through the use of memory-mapping. This can be done in Python by loading the input data from disk via *numpy*.*memmap* instead of *numpy*.*fromfile*, applied either to the original matrix (in the case of dense data) or on the “data”, “indices”, and “indptr” arrays (in the case of CSR arrays).

Because these arrays are often traversed in predictable fashion, typically sequential, we can off-load to disk, running transparently on datasets which significantly exceed machine RAM even in compressed form.

We also use memory-mapping by default in localsearch++, as an array of size (*k, n*_*points*_) may exceed available memory, and its sequential access patterns are convenient for memory-mapped data.

#### Algorithm 1

Generic Algorithm for Sparsity-Preserving distance computations given a prior adjustment *β*

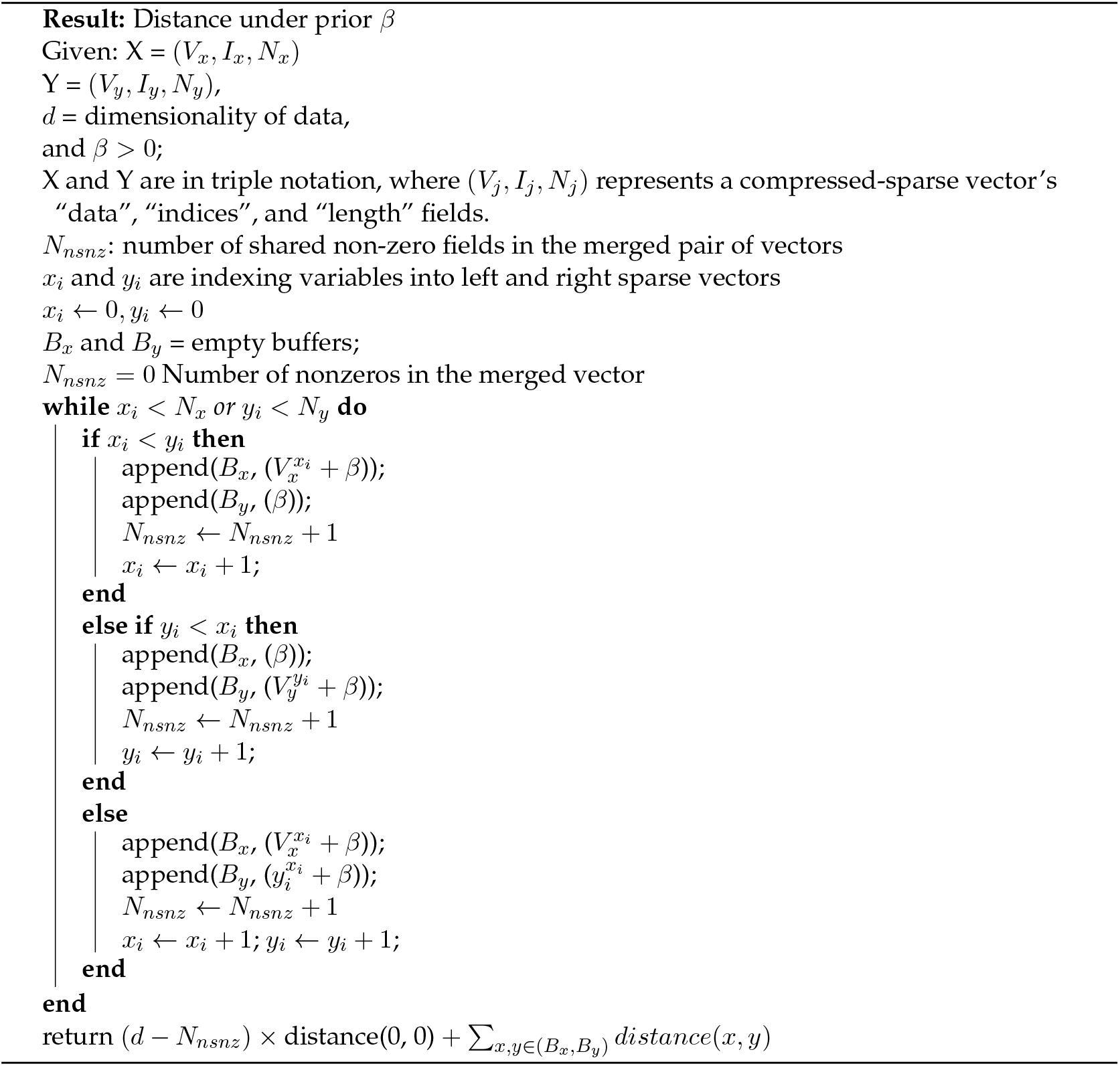

## 5 Acknowledgments

We thank Daniel Lemire and Wenzel Jakob for their fast SIMD Polynomial Congruential Generator Pseudorandom Number Generators.

Part of this research project was conducted using computational resources at the Maryland Advanced Research Computing Center (MARCC).

